# Concurrent infection of human brain with multiple species of Lyme disease spirochetes

**DOI:** 10.1101/2023.04.27.538619

**Authors:** Maryna Golovchenko, Jakub Opelka, Marie Vancova, Hana Sehadova, Veronika Kraliková, Martin Dobias, Milan Raska, Michal Krupka, Kristyna Sloupenská, Natalie Rudenko

**Author notes:** Address correspondence to Natalie Rudenko. Maryna Golovchenko and Natalie Rudenko contributed equally to this work.

## Abstract

Lyme disease (LD) spirochetes are well known to be able to disseminate into the tissues of infected hosts, including humans. The diverse strategies used by spirochetes to avoid the host immune system and persist in the host include active immune suppression, induction of immune tolerance, phase and antigenic variation, intracellular seclusion, and, importantly, incursion into immune privileged sites such as the brain. Invasion of immune privileged sites, like the brain allows the spirochetes not only escape from the host immune system but also can reduce the efficacy of antibiotic therapy.

Here we present a case of the detection of DNA of spirochetes from *Borrelia burgdorferi* sensu lato complex from multiple loci of LD patient’s post-mortem brain. The presence of co-infection with *Borrelia burgdorferi* sensu stricto and *Borrelia garinii* in LD patient’s brain was confirmed by PCR. The presence of atypical spirochete morphology was confirmed by immunohistochemistry of the brain samples and also in tissues of experimental mice, infected with *Borrelia* by simultaneous injection of spirochetes subcutaneously and intraperitoneally. Even though both spirochete species were simultaneously present in brain, the brain regions where the two species were detected were different and non-overlapping.

## Introduction

Lyme disease (LD) is a multi-system disorder with a diverse spectrum of clinical manifestations. It is caused by spirochetes of the *Borrelia burgdorferi* sensu lato (s. l.) complex. The initial stage of infection is characterized by flu-like symptoms (malaise, fatigue, headache, arthralgias, myalgias, fever, and regional lymphadenopathy) and/or, sometimes, skin rash (various forms of erythema migrans) developing within a few weeks after the tick bites. If the causative agent is not eliminated, it will further disseminate to the secondary sites of infection, leading to multiple subacute and persistent inflammatory pathologies, particularly affecting the central nervous system (CNS), joints or heart (1). Symptoms of the secondary stage of infection vary and may disappear after days to months, or continue, as the disease transition to the late stage of persistent disease with various signs and symptoms including fatigue, sleep disruption, cognitive deficits, arthralgia, myalgia, and headache. Even though antibiotic treatments in vast majority of infected patients result in full recovery, some patients suffer from long lasting neurological and psychological manifestations. Antibiotic therapy at late stage disease exhibits unpredictable response in resolution of symptoms (2-5).

For those who received standard antibiotic treatment, problems persisting more than six months after antibiotic treatment are termed, sometime, Post-Treatment Lyme Disease Syndrome (PTLDS) (6). The etiology of these syndromes is unknown, but several hypotheses have been discussed: *Borrelia* persistence, induction of immune disbalance leading to inflammation or autoimmunity, or disrupted central neural pathways leading to central sensitization, among others. Whether it depends on or is independent of microbial persistence it remains a topic of debate (7-8), however, the ability of LD spirochetes to colonize the multiple host tissues has been confirmed (9). Dermis is the first tissue that spirochetes colonize after the tick bite. At this point host immune system controls the pathogen burden in the tissue (10). Colonization of distant tissues involves the spirochete dissemination from the dermis, which is mediated by differential regulation of virulence determinants of *Borrelia* (11), that support the migration of pathogens through the endothelial and blood-brain barriers (12). Once established in immune privileged site, the pathogen is capable of triggering the local inflammation but is safe from being cleared by the host immune system and antibiotics, that can’t penetrate the blood-brain barrier (13). Survival of spirochetes despite antibiotic treatments, leading to the establishment of chronic LD has been clearly shown in animal studies (2, 14-17).

More spirochete species from *Borrelia burgdorferi* sensu lato complex are responsible for LD in Europe than in North America (18). The main burden of human cases, approximately 60%, is linked to *Borrelia afzelii.* The most common manifestation of *B. afzelii* infection is a skin lesion, erythema migrans (19). The second, most represented European *Borrelia* genospecies, *Borrelia garinii*, causes the Lyme neuroborreliosis (LNB), affecting both the central and peripheral nervous system (20) and up to 15% of LD patients suffer of LNB (13, 20). Intracellular localization of LD spirochetes in neurons and glial cells have been confirmed both, *in vitro* and *in vivo* (7, 21-22). Evidence of *Borrelia* persistence in the brains of chronic LNB patients is very limited; nevertheless, the development of dementia, cortical atrophy or amyloid deposition in some cases has been confirmed (7, 21). Invading neurons and glial cells, LD spirochetes can trigger progressive cell death or cause cell dysfunction (23). *Borrelia burgdorferi* sensu stricto (s.s.), is the major cause of LD in North America, however, its impact in European LD is under-appreciated (24-25). The most frequent manifestation of *B. burgdorferi* s.s. induced neuroborreliosis in the United States is lymphocytic meningitis whereas European *B. garinii –* induced LNB, in the majority of cases, is diagnosed as subacute painful meningoradiculitis or cranial nerve palsy, a uncommon manifestation of LNB in the USA (26). Such differences in clinical manifestations of LNB, caused by different spirochete species might be based on different mechanisms of dissemination of the bacterial pathogen into the nervous system, different capabilities of individual species to cross the blood-brain barrier (BBB) by either by transcellular or paracellular penetration (27), or different diagnostic protocols. Until recently, no reliable system for the detection of persistent infection exists. A single study on non-human primates showed that persistent forms of spirochetes that survived antibiotic treatment remain metabolically active (28). Because access to samples for study the persistence in humans is extremely difficult, studies rely either on findings generated in the animal studies or on analysis of post-mortem human originated specimens.

Here we report the case of the patient who, after being infected by *Borrelia* and treated with antibiotics continuously, progressed toward neurologic/psychiatric symptoms within the subsequent 13 years. After this period the patient underwent repeated serological testing with borderline positivity for *Borrelia* infection followed by prescription of several antibiotics, which provided no clinical improvements, followed by hospitalization at psychiatric clinics. Several months after releasing from the clinic, the patient committed suicide providing written consent for analyzing his brain for *Borrelia* presence.

## Materials and methods

### Ethical statement

Samples that were analyzed in this study originate from a young adult male who committed suicide in August 2019. The processing of post-mortem samples was performed based on informed consent provided by him before suicide in the form of a letter which was accepted by the Ethical Committee of University Hospital Olomouc, 102/18 (NV19-05-00191).

### CSF and blood collection and analysis

The autopsy was performed at the Institute of Forensic Medicine and Medical Law University Hospital Olomouc and Faculty of Medicine and Dentistry, Palacky University Olomouc, Czech Republic two days after the suicide. Peripheral blood and cerebrospinal fluid (CSF) samples were collected in a volume of 2 ml. The CSF was markedly stained by the presence of blood. Part of the CSF was aseptically removed for cultivation of spirochetes (Supplementary Materials and Methods). The remaining volume of CSF was centrifuged (2.000xg, 10 min, 4°C) to remove cells, and the supernatant was collected and stored at -80°C. The blood was centrifuged (2.000xg, 10 min, 4°C), clarified serum was collected, transferred to a new tube and stored at -80°C. Serological tests were performed by standard protocols using Anti-Borrelia EUROLINE-RN-AT, EUROLINE Autoimmune Inflammatory Myopathies 16 Ag (IgG) and EUROLINE ANA Profile 3 plus DSF70 (IgG) (EUROIMMUN, Luebeck, Germany) blot diagnostic kit with evaluation by a flatbed scanner and software EUROLineScan Software 3.4 (EUROIMMUN, Lübeck, Germany).

### Brain tissue collection (post-mortem)

Brain tissue samples of about 9 cm^3^ from seven different parts of the suicides’ brain were collected post-mortem by a certified pathologists in the Institute of Forensic Medicine and Medical Law, University Hospital Olomouc. Brain samples were collected from: 1 - temporal lobe (right), 2 – choroid plexus (left), 3 - occipital lobe (left), 4 - frontal lobe (left), 5 - parietal lobe (right), 6 - basal ganglia (right), and 7 - cerebellum (right). All tissue samples were divided into three parts. One part, that was used for cultivation of potentially live spirochetes. This sample was immediately aseptically transferred to BSK medium supplemented with 6% rabbit serum and antibiotics (as described above). The second part was used for PCR analyses. This portion was frozen at -80°C until use. The third part, that was used for immunohistochemical analyses. This part was fixed in 10% buffered formalin (4% paraformaldehyde) and stored at 4°C.

Based on PCR results, 5 samples of 125 mm^3^ in size were subsequently taken from the fixed autopsied occipital lobe sample for immunohistochemical staining (for the sample position within the occipital lobe autopsy is shown in Figure 1.

**Figure 1.**
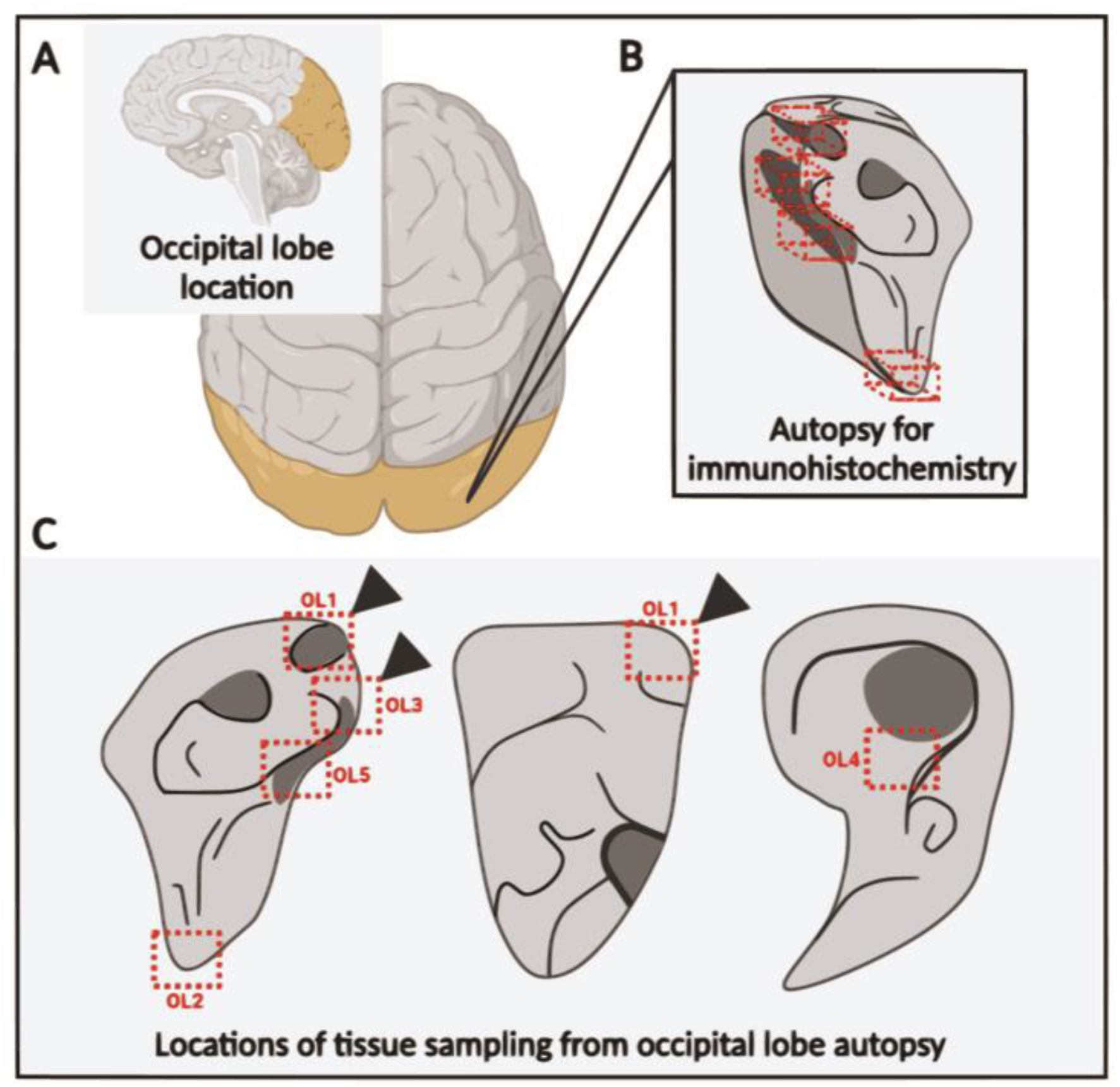
The scheme of the occipital lobe autopsy with description of sampling for the immunohistochemistry detection. (A) Location of the occipital lobe (yellow) in the human brain. (B) The model of the collected autopsy with dimensions (6×4×2 cm, approximately). (C) The red boxes and cubes indicate five screening sites of about 125 mm^3^ used for the immunohistochemical investigation. Arrowheads point to the locations where a positive signal was found.

### Analysis of total DNA from human brain tissues: polymerase chain reactions (PCR)

The DNA purification steps (Supplementary Material and Methods), PCR and post-amplification analyses were all performed in separate areas with all precautions against contamination. The presence of *Borrelia burgdorferi* s. l. DNA in the samples was assessed by PCR amplification of partial genes encoding outer surface protein C (OspC) and flagellin, followed by amplification of 8 housekeeping genes according to the previously described MLST protocol (29). To reduce the inhibition of the reaction from the excess of human DNA in the template DNA, the PCR reactions were conducted as nested PCRs under conditions previously used with human samples (30-31). The partial *ospC* and *flagellin* genes were amplified by nested PCR using the previously described primers (Supplementary Table S1) and conditions of the reaction as follows: 30 cycles of denaturation 95 °C for 30 sec, annealing 50°C and 52°C for 30 sec for the external and internal round of PCR respectively, and extension 72 °C for 30 sec (32-33). Two steps amplification of eight housekeeping genes included: *clpA* (BB0369), *clpX* (BB0612), *pyrG* (BB0575), *uvrA* (BB0837), *pepX* (BB0627), *recG* (BB0581), *rplB* (BB0481, seminested PCR), and *nifS* (BB0084, seminested PCR) (Supplementary Table S1). The PCR conditions for the housekeeping genes, except for *recG,* were as follows: initial denaturation 95°C for 15 min, cyclic denaturation 94°C for 30 sec, annealing temperature from 55°C to 48°C, (touchdown PCR, decreasing 1°C each cycle) for 30 sec, and the extension step of 72°C for 60 sec. An additional 20 cycles were run using denaturation temperature 94°C for 30 sec, annealing temperature of 48°C for 30 sec, and extension at 72°C for 60 sec. After a final extension step at 72 °C for 5 min, the samples were kept at 14°C until the second (nested) PCR. The conditions for the second PCR was as follows: 95°C for 7 min, followed by 35 cycles of [denaturation 94°C for 30 sec, annealing 50°C for 30 sec, extension 72°C for 60 sec]. After a final extension step for 5 min at 72°C, the samples were kept at 14°C.

For *recG*, the PCR conditions for the first set of cycles consists of initial denaturation 95°C for 15 min, followed by 30 cycles of denaturation at 94°C for 30 sec, annealing 55°C for 30 sec, extension 72°C for 60 sec, and final extension at 72°C for 5 min. The conditions for the second set of cycles were identical for all primers used.

The PCR reactions were carried out in a final volume of 20 μl using 2x HotStarTaq Plus Master Mix (Qiagen). Amplicons were visualized by electrophoresis in a 1.5 % agarose gel (1 × TAE, pH 8.0). In all cases, a reaction mix with water instead of a DNA template was used as the negative control. *Borrelia carolinensis* DNA was used as a positive control in all PCR reactions.

### Immunohistochemical detection of Borrelia in paraplast section of human brain autopsy and infected mouse tissues

Mice were infected with *B. burgdorferi* s.s. administered simultaneously by intradermal and intraperitoneal routes (Supplementary Material and Methods). The fixed autopsied occipital lobe samples were washed several times in phosphate-buffered saline (PBS). Organs dissected from euthanized mice were submersed in a fixative of saturated picric acid, 4% formaldehyde and 2.3% of copper acetate supplemented with mercuric chloride (Bouin-Hollande solution) (34) overnight at 4 °C. The fixative was then thoroughly washed with 70% ethanol. Standard techniques were used for both human autopsies and mouse tissue samples including, dehydration, embedding in paraplast, sectioning to 10 μm, deparaffinization and rehydration. The sections were treated with Lugol’s iodine followed by 7.5% solution of sodium thiosulphate to remove residuals of heavy metal ions, and then washed in distilled water and PBS supplemented with 0.3% Tween 20 (PBS-Tw). The nonspecific binding sites were blocked with 5% normal goat serum in PBS-Tw (blocking solution) for 30 min at room temperature (RT). Incubation with rabbit polyclonal *B. burgdorferi* antibody (Invitrogen, USA, specific to pool of *B. burgdorferi* s. l. complex proteins) diluted 1:200 in the blocking solution was done in a humidified chamber overnight at 4°C followed by washing the samples by thorough rinsing with PBS-Tw (three times for 10 min at RT). For enzymatic staining, the sections were further incubated in the cross-adsorbed horseradish peroxidase-labeled goat anti-rabbit secondary antibody (Invitrogen, USA) diluted 1:500 in the blocking solution for 90 minutes in RT, washed in PBS-Tw (three times for 10 min at RT), in 0.05M Tris-HCl pH 7.5 (for 10 min at RT) and stained in 10% 3,3’ diaminobenzidine in 0.05M Tris-HCl pH 7.5 with 0.005% H_2_O_2_ for 10 min in RT. The reaction was stopped by rinsing in distilled water, dehydrated and mounted in DPX mounting medium (Fluka, Switzerland). The samples were investigated under BX51 microscope equipped with DP80 CCD camera and cellSens software (Olympus, Tokyo, Japan) and the images were reconstructed by stitching of several Z stacks series. The further 3D image analyzes were performed in FIJI ImageJ software (35) using Interative Deconvolution 3D plugin.

For fluorescence staining, the samples treated with primary antibody and washed in PBS-Tw were incubated with goat anti-rabbit IgG conjugated Alexa Fluor 488 (Life Technologies, USA) diluted 1:500 in the blocking solution for 90 minutes in RT followed by rinsing with PBS-Tw (three times for 10 min at RT in the dark). The samples were dehydrated and mounted in DPX mounting medium (Fluka, Switzerland). The fluorescence signal was examined under the laser scanning confocal microscope FLUOVIEW™ FV3000 (Olympus, Japan) using the IMARIS software (Oxford Instrument, UK) for 3D reconstruction of the Z-stack series. Due to the autofluorescence of human brain autopsy samples, enzymatic detection of bound primary antibodies was preferentially used for the detection of *Borrelia* in these samples.

The specificity of the primary antibody to recognize both spiral and atypical forms of *B. burgdorferi* was verified by application of antibody on the paraplast sections of antibiotic-treated *B. burgdorferi* cultures mounted in agar (Supplementary Figure 1).

## Results

### Patient history and Borrelia serology

A male patient born in 1996, contacted a physician in 2004 after appearance of erythema migrans. He was diagnosed with LD and was treated with antibiotics (type and duration of treatment is unavailable), after which the patient suffered from neurological symptoms, mostly cognitive deficits such as “brain fog”, reduced psychomotor performance, and difficulties with concentration and processing of visual and auditory stimuli. In 2017 the patient was examined in the neurology department and had a positive *Borrelia* serology test, with both, IgG and IgM anti-borrelia antibodies in serum, but a negative PCR from a lumbar puncture. In December 2017, the patient was admitted to the Psychiatric Department of the University Hospital Olomouc. His therapy started with Zyprexa (Olanzapin) and was later replaced with Brintellix (Vortioxetin). On February 2018, anti-*Borrelia* antibodies were determined again in a private medical facility with borderline positivity for *Borrelia*-specific IgM and strong “++” positivity for IgG by ELISA. The ELISA results were confirmed by immunoblotting; the IgM immunoblot was positive for present OspC of *B. afzelii, B. garinii* and *B. burgdorferi* s.s. and the IgG immunoblot was borderline positive for VlsE of *B. afzelii, B. garinii* and *B. burgdorferi* s.s., and negative for p83, flagellin, BmpA, OspA, OspB, OspC and DbpA. Serological tests for *Chlamydia sp., Mycoplasma sp., Anaplasma phagocytophillum, Toxocara canis* and *Toxoplasma gondii* were negative. A PCR test for the presence of *Babesia* sp. in peripheral blood was negative. At the same private medical facility, patient was prescribed a combination of antimicrobials: Minocycline (3×100mg/day), Azithromycin 250 mg (3x/week), and Hydroxychloroquine (Plaquenil 200 mg, 1×1). In August 2018, the suspected hypocorticism was ruled out by biochemical laboratory examinations. In September 2018 the patient was hospitalized at the Department of Psychiatry University Hospital Olomouc with suspicion of a developing mental disorder. The patient was diagnosed with schizotypal and somatomorphic condition and was discharged three weeks later. In August 2019 the patient committed suicide. The patient’s body was dissected in the Institute of Forensic Medicine and Medical Law, University Hospital Olomouc 2 days after the death. The patient left behind the letter expressing the urgent demand to scientists to analyze his brain for presence of LD spirochetes. The letter was provided to the Ethical committee of University Hospital Olomouc and initiated this study.

### Post-mortem toxicology and microbiology

Toxicological post-mortem analysis confirmed the presence of following chemical agents in the blood: hydroxychloroquine, hydroxyzine (the active substance in the anxiolytic Atarax) and its metabolite cetirizine. In addition, a low concentration of azithromycin was detected in CSF. This confirms that the patient was taking the prescribed combination antibiotic therapy until shortly before death. Immunoblot analyses of post-mortem serum and CSF conducted at the Faculty of Medicine and Dentistry, Palacky University Olomouc, confirmed borderline IgG reactivity against VlsE of *B. garinii* and *B. burgdorferi* s.s. The reactions with VlsE antigens of *B. afzelii*, lipids of *B. garinii* and *B. afzelii*, p83, p41, p39, OspC, p58, p21, p20, p19 and p18 were negative. Using immunoblot kits for autoantibodies detection, weak IgG positivity against PL-7 antigen (threonyl-tRNA synthetase) and borderline reactions with SRP antigen and histones were detected. As a part of the microbiological analysis of post-mortem samples, the CSF and brain tissues were used for the cultivation of spirochetes in the BSK-H medium. After 2 months of incubation of seeded cultures under conditions described above, the presence of live bacteria were not detected in any culture and all samples were deemed to be culture-negative.

### PCR detection of Borrelia DNA in frozen brain tissue samples

To determine if *Borrelia* DNA could be detected in the brain, PCR analysis was performed on DNA from seven different parts of the human brain, samples of which had been frozen at -80°C upon collection. Nine genes (*flaB, ospC, clpA, clpX, nifS, rplB, pepX, pyrG* and *uvrA*) provided the amplicons of the expected size (Table S1), although the PCRs were not 100% successful for all tested brain loci (Table 1).

**Table 1.** Results of PCR amplification of *Borrelia* genes in different brain loci

| gene<br>sample | *fla B*<br>388 bp | *ospC*<br>617 bp | *clpA*<br>706 bp | *clpX*<br>721 bp | *nifS*<br>629 bp | *rplB*<br>720 bp | *pepX*<br>666 bp | *pyrG*<br>687 bp | *uvrA*<br>677 bp |
| --- | --- | --- | --- | --- | --- | --- | --- | --- | --- |
| 1- temporal lobe | *Bb* s.s. | *Bb* s.s. | + |  | + | + | + |  | + |
| 2- choroid plexus | *Bb* s.s. |  | + |  | + | + | + | + | + |
| 3- occipital lobe |  | *Bb* s.s. | + |  | + | + | + | + | + |
| 4- frontal lobe | *Bb* s.s. | *Bb* s.s. | + |  | + | + | + | + | + |
| 5- parietal lobe | *Bb* s.s. | *Bb* s.s. | + | + | + | + | + | + | + |
| 6- basal ganglia | *B. gar* | *B. gar* |  |  | + | + | + | + | + |
| 7- cerebellum | *B. gar* |  |  |  |  | + | + | + | + |

“+”- DNA fragment of expected size was obtained by PCR amplification

Sequence confirmation, followed by comparison with available databases, confirmed the presence of DNA of two spirochete species, *B. burgdorferi* s.s and *B. garinii*. The different *Borrelia* genospecies were differently distributed in brain samples: DNA of a single species, *B. burgdorferi* s.s., was detected in temporal right lobe, choroid plexus (left), occipital lobe (left), frontal lobe (left) and parietal lobe (right)), while the DNA of *B. garinii* was identified in the basal ganglia (right) and cerebellum (right). No PCR amplification showed the presence of more than one spirochete species in any sample.

BLASTN analysis of the *ospC* sequences confirmed that the *B. burgdorferi* s.s. strain carried *ospC* of type A, the most common *ospC* type that is widely distributed globally and is considered to be the most invasive type. The *B. garinii ospC* sequences detected in the brain samples were identical to strains widely distributed in Eurasia.

### Occipital lobe tissue exhibited structures consistent with Borrelia

Immunohistochemical investigation of paraffin sections of the patient occipital lobe samples (Figure 1) revealed the presence of structures reactive with *Borrelia*-specific rabbit polyclonal antibodies (Figure 2). The size and morphology of these structures resembled *Borrelia* cells with the atypical morphology detected from cultured spirochetes treated with doxycycline or amoxicillin, used as an *in vitro* test of antibody specificity (see Supplementary Material and Methods and Supplementary Figure 1). In *Borrelia* populations treated by both low (50 μg/ml) and high (100 μg/ml) concentrations of either doxycycline or amoxicillin, the antibody stained both spiral and atypical *B. burgdorferi* forms (Supplementary Figure 1 A, C, E, G, I). The presence of both forms of *Borrelia* in the culture was verified using transmission electron microscopy (Supplementary Figure 1 B, D, F, H, J). The structures detected in the samples of the patient occipital lobe had a diameter of about 1-10 μm, and presumed protoplasmic cylinders ranged between 0.2 - 0.4 µm in diameter. These structures were detected with an estimated frequency of 0.16 - 0.3 per 1 mm^3^ of tissue typically located near the capillaries (Figure 2 A, B). These findings were in accordance with the observation of tissue samples from mice artificially infected with *B. burgdorferi* s.s. (Figure 3), where structures corresponding to the atypical forms of *Borrelia* were detected. Out of all tested mouse tissues, these *Borrelia*-like structures were mainly found in samples from bladders (Fig. 3 A, B) and knee joints (Fig. 3 C, D).

**Figure 2.**
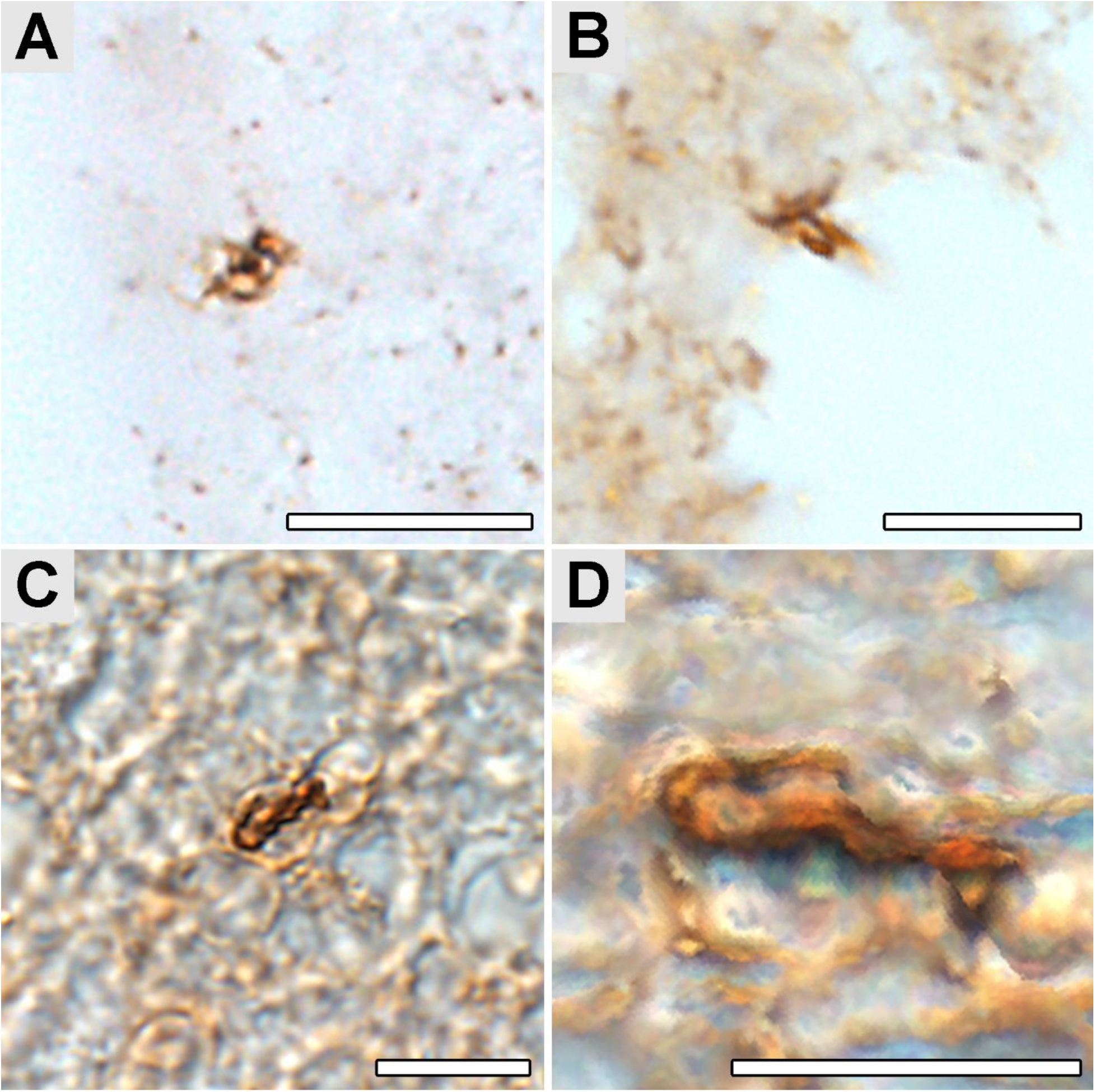
*Borrelia*-like structures in the paraplast section of a sample from the patient’s occipital lobe. Anti-*B. burgdorferi* polyclonal antibody visualized by the horseradish peroxidase-conjugated secondary antibody was used for the detection. (A-C) Images from a light microscope showing structures ressembling atypical forms of *Borrelia*. (D) The image C edited by image analysis in the FIJI software. Scale bar: 10μm.

**Figure 3.**
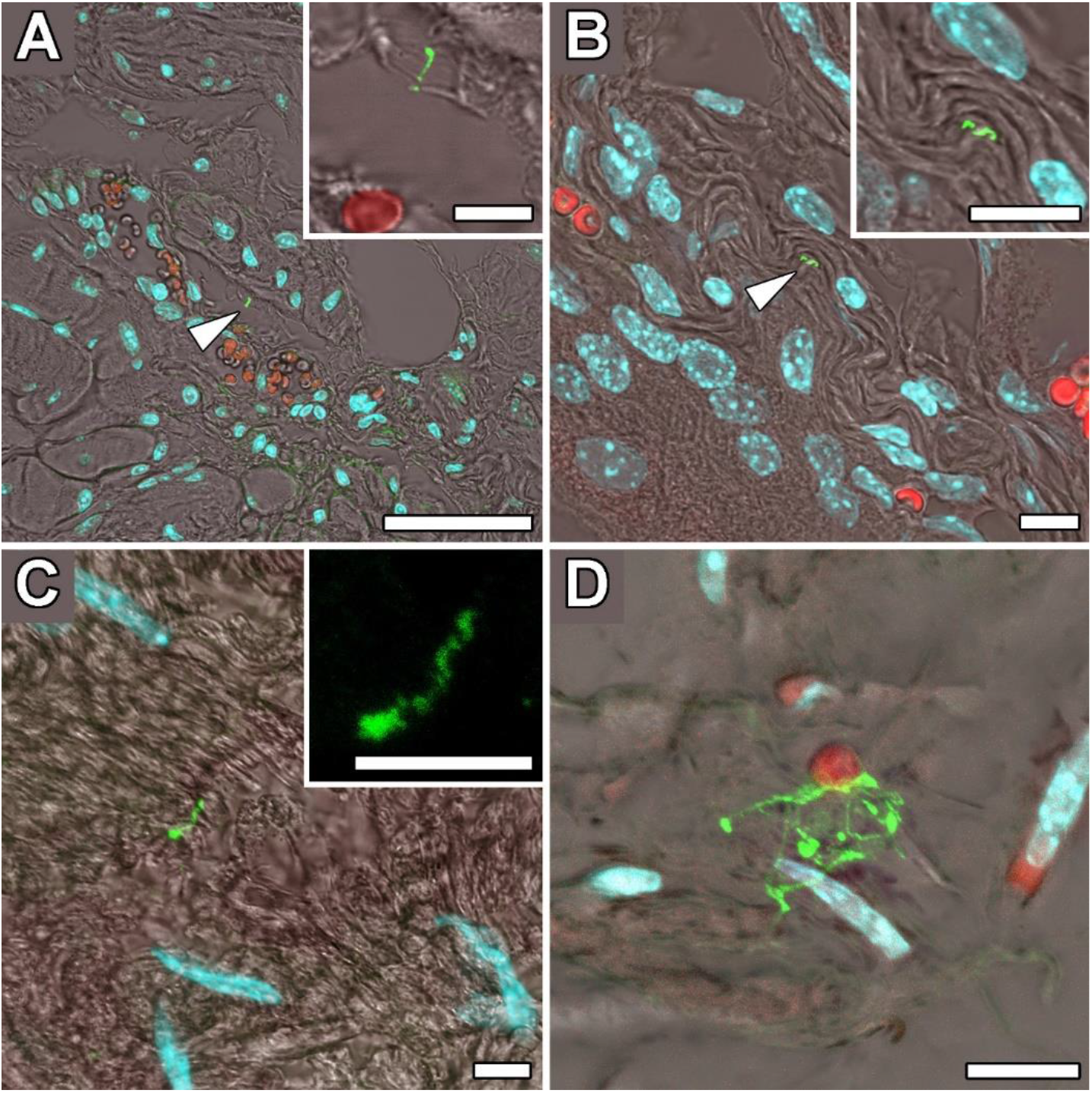
Laser scanning confocal images of detected *B. burgdorferi* sensu stricto in paraplast tissues sections of the C3H/HeN mouse line treated with doxycycline or amoxicillin. (A, B) Persistent form of *Borrelia* in the bladder (arrowheads). (C, D) Persistent form of *Borrelia* in the knee joint. Insets show close-up views of atypical forms of *Borrelia*. *Borrelia* detected by anti-*B. burgdorferi* polyclonal antibody and visualized by fluorescently labeled secondary antibody (Alexa Fluor 488), nuclei stained by DAPI (in blue), and autofluorescence of erythrocytes (in red). Scale bar: A 100 μm, B, C, D, and insets 10 μm.

## Discussion

Detection of *Borrelia* in multiple organs of infected animals, including humans, demonstrates the ability of spirochetes to disseminate into the secondary sites of infection (to review see 7 [citations 17-49]). Detection of intact spirochetes at the sites of secondary infection in both laboratory animals and humans after even aggressive antibiotic treatments further demonstrates the ability of the spirochetes to persist (13, to review see 7 [citations 50-85]). These previous findings are consistent with our finding of spirochetes in a human brain despite extended antibiotic treatment. Earlier presented detection of intact spirochetes in autopsy brains specimens of the human after extended treatments and these with diverse history of disease manifestations, from widely recognized neurocognitive disorder, anxiety, depression, memory loss to brain atrophy and progressive dementia (21, 36-40), might show that persistent *Borrelia* infection can lead to persistent disorder of central nervous system (CNS) (13, 41-42). Neurotropic nature of LB spirochetes secures its survival in the CNS (43) for an extended period of time in alternative morphological forms (44-46), including antibiotic-resistant biofilms (47). In general, dissemination of spirochetes occurs through the bloodstream where the spirochetes remain for a short period of time, heading finally to the extracellular matrix of varied internal organs, where they retain protection from a host immune system or antibiotics (11, 48-51). However, LB spirochetes were found to be present as well intracellularly in cardiac myocytes, endothelial or synovial cells (52-54). Dissemination of spirochetes into the CNS is thought to occur via passage along the peripheral nerves (23). Based on the distinct clinical manifestations of LNB in Europe and North America, caused by *Borrelia garinii* and *B. burgdorferi* s.s., respectively, the mechanisms of spirochete dissemination into the CNS might be species-dependent and rely on *Borrelia* ability to cross the BBB (26-27, 55).

Our results confirmed the presence of two species of spirochetes from *Borrelia burgdorferi* sensu lato complex, *Borrelia garinii* and *Borrelia burgdorferi* s.s., in different areas of the human brain. Importantly, the DNAs of two spirochete species were detected in distinct areas of the brain; in no case did we find infection with both in the same brain region. Multiple repeated PCR amplifications with different sets of primers, followed by sequence confirmation of the *Borrelia* species, always provided clear and definite results indicating the presence of only one species. Based on the analysis of a brain from only a single individual, it is not yet possible to conclude whether this finding represents biological restriction in brain colonization or is just a coincidence. The mechanism of crossing the BBB by spirochetes is not understood yet as a good animal model has not yet been established. Recently used inbred mice as a model have shown colonization of dura mater during acute and late spirochete infection (56-57). The development of LNB depends on the ability of spirochetes to cross the BBB. This invasion can occur via breaching either physical (tight junctions) or metabolic (enzymes, transport systems) barriers (58). The role of the BBB in neurodegenerative and neuropsychiatric disorders is crucial; its failure plays an important role in the pathogenesis of many diseases of the CNS that are caused by bacteria or protozoa (59-61). Recent studies show that, in the case of some neurocognitive disorders that lead to the development of dementia, as well as during normal aging, the “leakage” of the BBB is increased (62-63). Whether *Borrelia* enter the brain by direct transmigration or are carried across the BBB hidden inside non-phagocyting leukocytes, using the “trojan horse” mechanism as in the case of other pathogens, is not yet known (60-61). It is tempting to speculate that some kind of competition might occur between the spirochete species in the process of crossing the BBB, and this might be the basis of our observations. A recent study by Adams and colleagues (27) showed that different species of LD spirochetes are able to enter BBB-organoids with different rates of success. The spirochetes that successfully invaded the organoids remained viable inside the BBB-organoids, initiating the loss of tight junction and changes in the organoids gross morphology and integrity (27). To our knowledge, this is the first study that confirms the presence of *B. garinii* and *B. burgdorferi* s.s. structures in the human brain with strict separation of invaded brain loci. The question of what defines the distribution of spirochete species in brain tissues remains unanswered but is one of increasing importance as the number of human *Borrelia* infections increase.

The attribution of persistent symptoms to Lyme disease and potential *Borrelia* persistence is limited by several factors including presence of comorbid disease with overlapping symptoms or reinfection with *Borrelia*. For example, in the case of symptoms such as fatigue, headache, sleep disruption, cognitive malfunctions such as feeling of poor concentration, confusion, slower thinking, forgetfulness, lost words or mental fatigue, sometimes identified by patients as a “brain fog”, it is difficult to exclude the contribution of endogenous psychiatric etiology even in the case of positive laboratory proofs of previous *Borrelia* exposure. Nevertheless, the important fact remains that several months of combined antimicrobial treatment in the presented case did not lead to an improvement in this patient’s condition. Human LNB is an example of a very complex disorder. Big controversy exist both in diagnosis and treatment of this disease, especially when it comes to persistent infection, chronic illness, PTLDS or the course of patients treatment with such conditions. It remains an open question whether long-term antimicrobial treatment may have contributed to the progression of neurological symptoms in reported patient. Two of the drugs he used are suspected to have neurological or neuropsychiatric side effects, namely minocycline (64-65) and hydroxychloroquine (66-68). Either long-term or repeated antibiotic therapy of PTLDS also carries a number of other risks, including development of significant, even life-threatening disorders such as necrotising enterocolitis or systemic candidiasis (69-70). Both can affect as well the finding and treating of other possible causes of the symptoms (71-72).

Since the recognition of Lyme disease, thousands of reports have been published, but the optimal therapy is still a matter of debate (73). The complexity of the pathophysiology of the causative agent of LD and the clinical uncertainty surrounding Lyme or other tick-borne diseases make the choice of treatment not straightforward. Recognized persistent bacterial infections may require prolonged antibiotic therapy and seem reasonable and justifiable in some situations when considering patients with persistent LD symptoms (73).

## Acknowledgments

This research was funded by grant NV19-05-00191 from Ministry of Health of the Czech Republic. Partial analysis of the samples was supported by project RVO 60077344 of the Biology Centre CAS, Institute of Entomology. We acknowledge the BC CAS core facility LEM supported by MEYS CR (LM2023050 Czech-BioImaging and OP VVV CZ.02.1.01/0.0/0.0/18_046/0016045).

## Institutional Review Board Statement

The study was conducted according to the guidelines of the Declaration of Helsinky, and approved by the Institutional Ethics Committee of University Hospital Olomouc (reference number 102/18 from June 2018). The protocol of the study, including the informed consent of the patients, was approved by the Ethics Committee of the Olomouc University Hospital (reference number 102/18 of June 2018).

## Conflicts of Interest

The authors declare no conflict of interest.

## Supplementary Material and Methods

### Induction of formation of atypical morphologies of Borrelia by antibiotics

Spirochetes of *B. burgdorferi* s.s. strain NE-5264 were grown in modified Kelly-Pettenkofer medium (MKP) (74). The cultures were incubated at 33°C until cell density reached at least 10^6^ spirochetes per milliliter. The absence of contamination and the viability of spirochetes was verified by microscopy. The spirochete concentration was determined using a Petroff-Hausser counting chamber. Five sterile DNase-free Eppendorf tubes with spirochete cultures were treated with antibiotics doxycycline or amoxicillin (Sigma-Aldrich, USA) regularly used in the treatment of LD. Each antibiotic was applied at two concentrations: 50 μg/ml and 100 μg/ml. An untreated culture was used as a positive control. After 14 days of antibiotic treatment, the spirochetes (7.5 × 10^7^) were washed in 0.1 M HEPES, pelleted by centrifugation (820 × g, 10 min), fixed in 4% formaldehyde with 0.1% glutaraldehyde in 0.1M HEPES for 1h at RT and immediately transferred to freshly prepared 2% agar for processing of paraplast sections.

### Infection of laboratory mice (control)

Susceptible to *Borrelia* mice C3H/HeN genotype were used as laboratory animal model for control experiments. Six weeks old female mice (Jackson Laboratory, Germany) were infected by simultaneous subcutaneous and intraperitoneal injections of 10^4^ replicating spirochetes in 100 μl of MKP medium per mouse.

### Immunohistochemical detection of cultured Borrelia

Immunodetection of *Borrelia* on paraplast sections of cultured spirochetes was performed using the same protocol as for human brain tissue (above). A specifically bound primary antibody was detected by incubation with the goat anti-rabbit IgG conjugated Alexa Fluor 488 secondary antibody (Life Technologies, USA), diluted 1: 500 in the blocking solution, for 90 minutes at RT in dark.

### DNA extraction

For total DNA purification from all collected samples, the DNeasy Blood and Tissue kit (Qiagen, Germany) were used. To increase DNA yield and so the possibility of capture of spirochete DNA in the sample, the entire frozen tissue was weighted, homogenized in liquid nitrogen and then subsequently processed according to the manufacturer’s protocol.

### Sequencing

All amplicons of the expected sizes were excised from agarose gels, purified using QIAquick PCR Purification Kit (QIAGEN, Germany) and sequenced in both directions using the same primers used for amplification. Sequence analysis was performed commercially by SEQme s.r.o. (Czech Republic) and the sequences were compared to those available in the NCBI GenBan database using Basic Local Alignment Tool (BLASTn) analysis.

### Cultivation of Borrelia from CSF samples

Five hundred microliters of CSF were transferred to a 5 ml of Barbour-Stoner-Kelly culture medium (BioConcept, Switzerland) supplemented with 6% rabbit serum (Merck, USA) and antibiotics phosphomycin, polymyxin and rifampicin (100x concentrated solution, HiMedia, India diluted 1:100). Seeded cultures were kept at +33°C for two months with regular checks by dark-field microscopy, starting from day 10 after culture initiation.

### Transmission electron microscopy

Spirochetes were fixed in 2.5% glutaraldehyde in 0.1M PBS for 1h at RT. Cells were washed three times in 0.1 M PBS with 4% glucose, embedded into 2% of agar, and postfixed in 2% OsO_4_ for 1h at RT. After washing, samples were dehydrated stepwise using a graded acetone series (30-50-70-80-90-95%, v/v) for 15 min at each step and transferred to absolute acetone for 15 min. Samples were infiltrated in 2:1, 1:1, and 1:2 mixtures from acetone/stock resin solutions (1h/each step) and finally in two changes of Poly/Bed 812 resin (Polysciences Inc., USA) before embedding and polymerization. Ultrathin sections were stained in saturated ethanolic uranyl acetate and lead citrate before imaging in JEM-1400 Flash TEM (JEOL Ltd.).

---

**Supplementary Figure 1.**
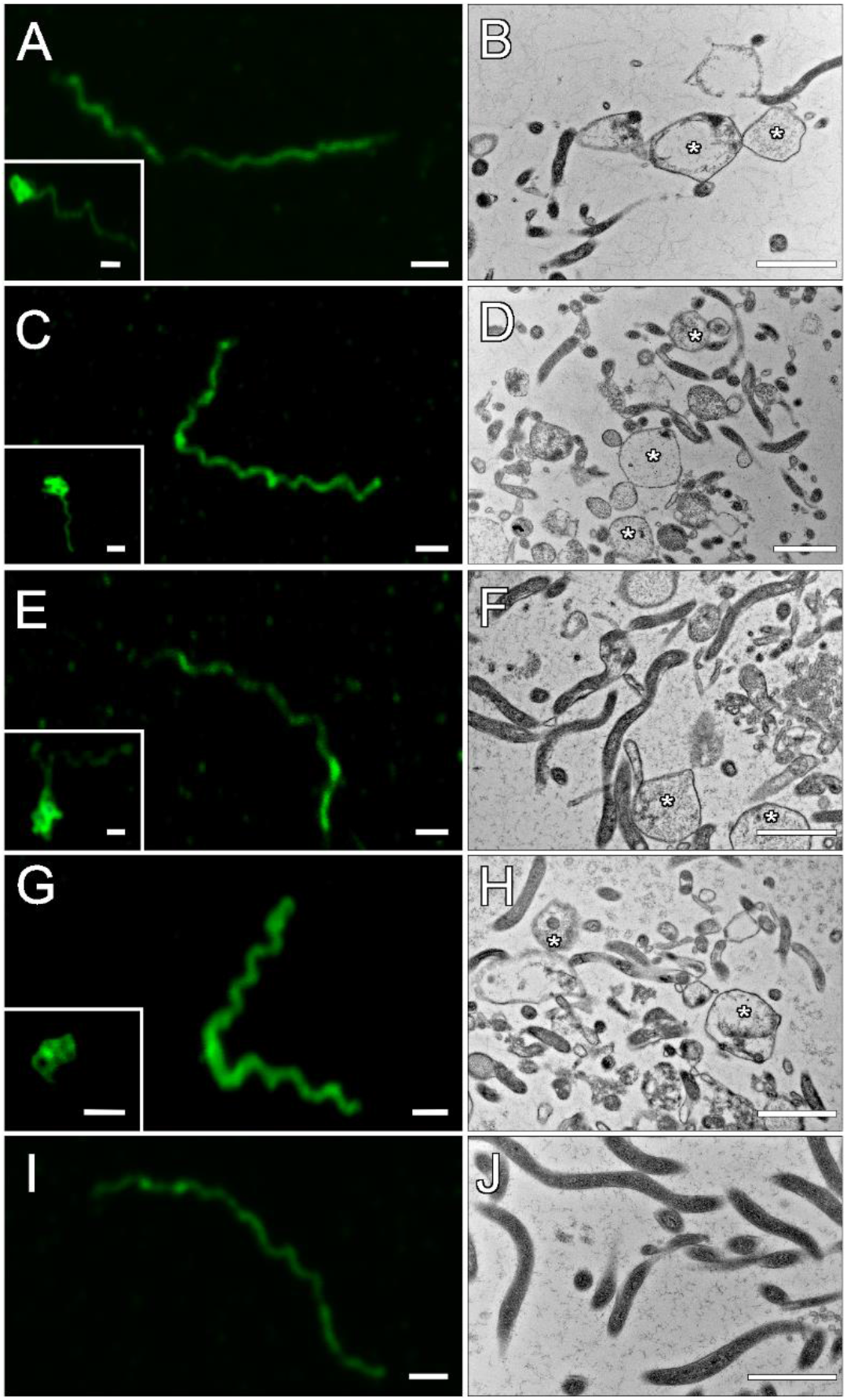
Spiral and atypical Borrelia forms detected after antibiotics treatment. Immunofluorescence and TEM images of *Borrelia burgdorferi* s.s. cultured with amoxycillin (A-D) and doxycycline (E-H) antibiotics at concentrations of 50 µg/ml (A,B, E,F) and 100 µg/ml (C,D,G,H). Both spiral and atypical morphological forms were observed in ATB-treated cultures in contrast to control (I, J). Scale: 1 μm.

**Supplementary Table S1.**
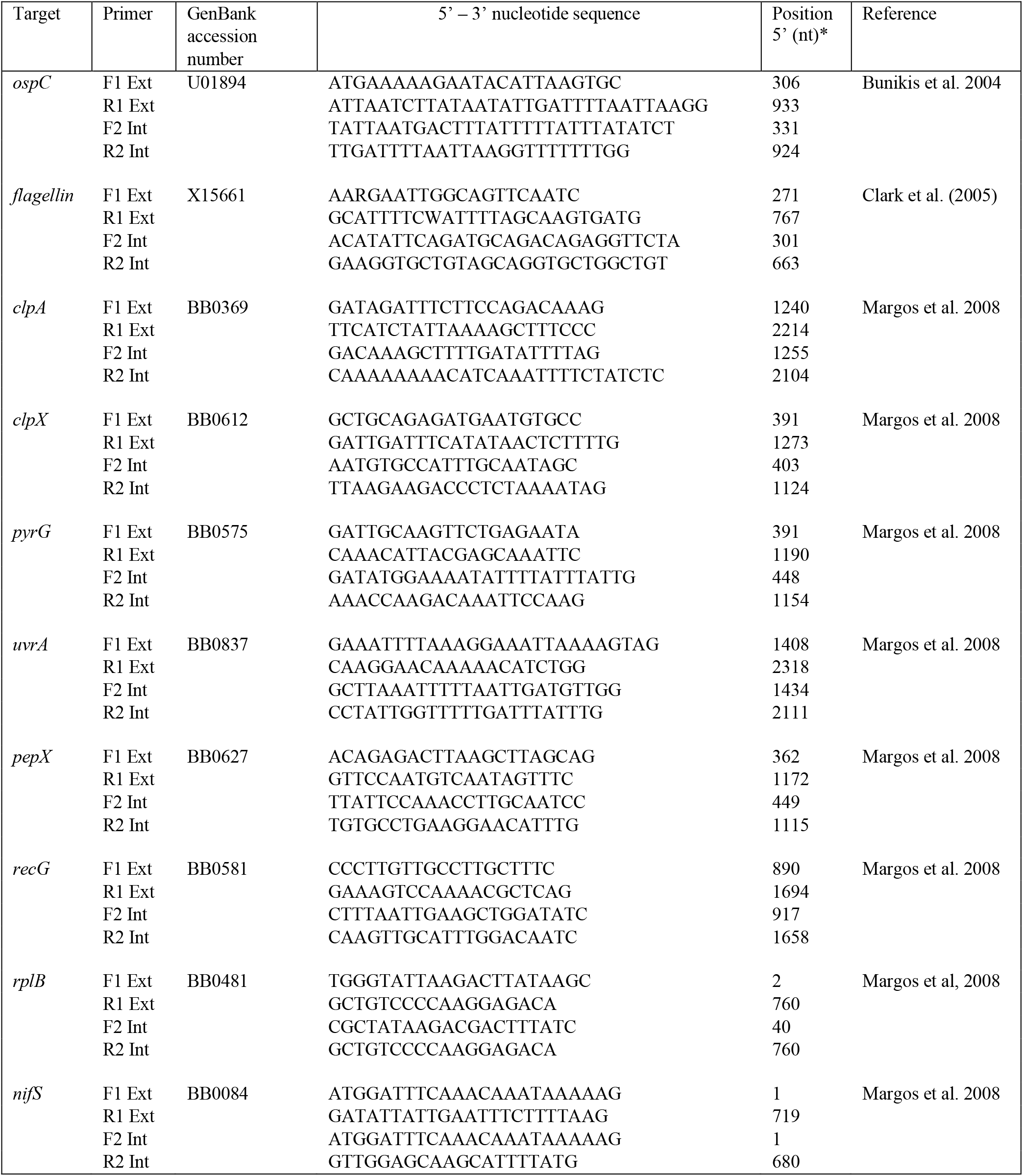

